# Simulated Microgravity Induces Cultivar-Specific Changes Affecting *Salmonella enterica* Ingression Independent of Stomatal Physiology

**DOI:** 10.64898/2026.05.13.724889

**Authors:** Tania A.M. Wiest, Harsh P. Bais

**Affiliations:** Department of Plant and Soil Sciences, University of Delaware, Newark, 19713, USA; AP Biopharma, University of Delaware, Newark, 19713, USA 18 19

**Keywords:** Microgravity, Spaceflight, Lettuce, Salmonella enterica, Stomata

## Abstract

Advances in NASA’s astrobiology program have demonstrated the feasibility of cultivating plants in space and in analog extraterrestrial habitats. In addition to abiotic stressors, plants grown in terrestrial and space-like environments are challenged by both phytopathogens and opportunistic human pathogens, with implications for plant productivity and human health. The persistence of human-associated pathogens in spacecraft and space stations raises significant concerns regarding food safety. The molecular, biochemical, and signaling mechanisms governing stomatal development and function under microgravity remain poorly understood. We employed an experimental system incorporating human pathogen *Salmonella enterica* and lettuce microgreens exposed to simulated microgravity through two-dimensional clinorotation to investigate plant innate immunity and stomatal development and function. We further evaluated four lettuce cultivars to determine whether genetic variation impacts these factors under simulated microgravity conditions. Our findings indicate that simulated microgravity significantly influences stomatal development and function, as evidenced by an increase in stomatal density and variable changes to stomatal aperture. Notably, cultivar-dependent variation in stomatal traits and responses to *Salmonella enterica* was observed under microgravity conditions. Although increased stomatal density was hypothesized to enhance pathogen ingression, internalization was more strongly predicted by cultivar selection and simulated microgravity; simulated microgravity increased ingression, with red pigmented cultivars having less pathogen than green cultivars. These results suggest that targeted selection of cultivars with favorable physiological traits may improve food safety and the viability of crop production systems in space environments. They also suggest that development and function of stomata may change in spaceflight conditions.

## Introduction

For deep space exploration to be possible, advanced habitation systems (AHS) must be self-sustaining, and growing crops during missions can address several key AHS needs, including water filtration, air quality improvement, and nutrition support (Kordyum and Hasenstein, 2021). Microgreens are gaining interest for space agriculture since they are ready to harvest within 1–3 weeks of planting, need minimal space and energy, and produce immune boosting phytonutrients that are difficult to encapsulate into supplements (Kyriacou et al., 2017). Microgreens are a concentrated source of a wide variety of phytonutrients that have a broad variety of functions. Most valuable to astronauts are antioxidants like glucosinolates, carotenoids, and phenols (Tallei et al., 2024). These non-essential nutrient compounds have extra value in space, as increased radiation exposure increases reactive oxygen species (ROS) generation (Zhao, 2024). While some ROS is essential for normal bodily function, an imbalance of ROS has been linked with increased aging and disease (Sánchez-Rodríguez and Mendoza-Núñez, 2019). Consumption of fresh produce and their antioxidants has been shown to combat cancer, improve cardiovascular health, protect the nervous system, reduce inflammation, and more (Tallei et al., 2024). Lettuce microgreens in particular have been shown to have 79% more ascorbic acid and phenols than mature counterparts, and red pigmented cultivars were more nutrient dense than green cultivars (Martínez-Ispizua et al., 2022). Combatting the many stresses of spaceflight on the human body is essential as these stressors could lead to more severe outbreaks of illness (Hummerick et al., 2021).

Outbreaks of foodborne illness are potentially a serious threat to space exploration missions. Some human pathogens, like *Salmonella enterica* serovar Typhimurium, have been shown to have improved virulence under both true and simulated microgravity (Sµg) (Higginson et al., 2016). Microbial analysis of space-grown lettuce has shown that a diverse and distinct microbiome is present on International Space Station (ISS) grown plants when compared to controls grown on Earth (Khodadad et al., 2020). Pre-flight analyses of food samples have found the presence of *S. enterica, Staphylococcus aureus*, *Enterobacter cloacae,* and *Cronobacter sakazakii.* Thus far, such tests have prevented the transport of most of these pathogens to the ISS (Colorado et al., 2021). If obligate human pathogens like *S. enterica* were to contaminate spaceflight systems, significant illness and death could occur. *S. aureus* has already been identified from space-grown Mizuna samples, though in densities unlikely to causes food poisoning (Bunchek et al., 2024). This could be because plants are not passive hosts to microorganisms; rather, they have an innate and inducible immunity to respond to plant and some human pathogens (Melotto et al., 2014). Plants recognize microbes through microbe-associated molecular patterns, including flagellin and lipopolysaccharides (Jones et al., 2024). Their ability to interact with diverse microbes stems from an immune system that differentiates between pathogenic and non-pathogenic plant–microbe interactions. Knowledge of how plants interact with opportunistic human pathogens such as *S. enterica* is still emerging. Multiple studies suggest that the persistence of human pathogens on or within lettuce tissues can be altered by lettuce cultivar and pathogen strain (Jacob and Melotto, 2020; Klerks et al., 2007; Yin et al., 2020). However, how the interaction between between pathogens, crop plants, and humans is modulated by spaceflight is critically understudied (Totsline et al., 2023).

Despite the wide evolutionary distance between plants and humans, numerous human pathogens—including *Salmonella enterica*, *Pseudomonas aeruginosa*, *Staphylococcus aureus*, *Escherichia coli* O157:H7, and *Shigella* spp.—exhibit cross-kingdom pathogenicity by infecting and/or colonizing both plant and animal hosts (Sobiczewski and Iakimova, 2022). Although typically associated with animal disease, these pathogens can persist on plant surfaces as epiphytes and within plant tissues as endophytes, where they occupy the apoplast—a sheltered and nutrient-rich environment (Melotto et al., 2014). To transition from an epiphytic to an endophytic lifestyle within the apoplast, bacteria must gain entry through natural openings or physical disruptions in plant tissues. These entry points include roots, wounds, trichomes, hydathodes, and stomata (Thomas et al., 2024). Among these, stomata represent a critical route for the internalization and persistence of pathogens (Saldaña et al., 2011). Stomata are microscopic pores covering above ground tissues that regulate gas exchange, flanked by two guard cells that can open and close the pore in response to abiotic and biotic environmental signals. Opening or closing stomata happens on a time-scale of minutes compared to hours or days like many other adaptations, making it one of the fastest ways for plants to adjust to their environment (Kollist et al., 2019). Foodborne pathogens can trigger stomatal immunity, the process of closing stomata after sensing a pathogen threat (Melotto et al., 2017). Furthermore, human pathogens can trigger, suppress, and evade stomatal and plant innate immunity using virulence factors similar to those employed by plant pathogens (Johnson et al., 2020). With some exceptions, human enteric pathogens are not considered true phytopathogens because they generally do not produce visible disease symptoms in plant hosts (Melotto et al., 2014). However, once inside a plant host, these pathogens can persist at medically relevant population levels within the apoplast (Schikora et al., 2012).

Successful colonization of the apoplast requires pathogens to evade or suppress the plant’s immune responses. Notably, some opportunistic human pathogens can bypass plant defenses, including stomatal immunity, in a manner analogous to plant pathogens (Garcia and Hirt, 2014). For example, *S. enterica* has been shown to reopen stomata in leafy green species under normal gravity conditions, facilitating entry into internal tissues (Johnson et al., 2020). Additionally, under Sµg conditions, lettuce plants maintain open stomata for longer periods, leading to increased *S. enterica* ingression (Totsline et al., 2024). Recent findings suggest that *S. enterica* may utilize bacterial-derived auxin biosynthesis to reopen stomata in *Arabidopsis*, thereby facilitating ingression (Fochs et al., 2025). However, whether *S. enterica* exploits auxin biosynthesis under microgravity, conditions remain unknown. Utilizing different lettuce cultivars to assess stomatal phenotypes, specifically stomatal density and stomatal aperture, under Sµg and *S. enterica* treatments may reveal the role of subtle genetic variation in improving food safety for space-based agriculture.

Despite being a critical route for the internalization and persistence of pathogens, not to mention an essential part of gas exchange regulation, how stomata develop and function in space is critically understudied. Most plants are hypostomatous, where all stomata are on the abaxial (bottom) surface. This trend towards hypostomaty could be to improve disease resistance against foliar pathogens that accumulate on the adaxial surface. This stomatal distribution could also conserve water, as the abaxial surface is cooler and has lower evapotranspiration rates compared to the top surface (Muir, 2015). However, many plants like lettuce, wheat, and *Arabidopsis* are amphistomatous with stomata on the adaxial (top) surface as well (Matkowski and Daszkowska-Golec, 2023). Emerging research suggests that amphistomatous plants differentially regulate top adaxial and abaxial surfaces (Jalakas et al., 2024), often placing more stomata on the abaxial compared to the adaxial surface, likely for the same reasons plants evolve hypostomaty (Muir, 2015). Plants tightly control the placement and density of these structures in order to maximize gas exchange while managing transpiration and water loss (Pillitteri and Torii, 2012). Stomata are also a point of entry for *S. enterica* to potentially enter a leaf, proliferating in an environment that sanitizing efforts cannot reach (Johnson et al., 2020). Previous work from by Totsline et al., (2024) found that Sµg did significantly alter stomatal defense against *S. enterica,* which led to deeper ingression under 2D clinorotation. How spaceflight affects stomatal patterning is largely unknown, along with how cultivar selection impacts stomatal patterning and defense in space.

In this study, we investigate how stomatal development and function in lettuce influence *Salmonella* ingression under microgravity conditions, and how cultivar selection impacts these changes. Our primary hypotheses were that 1) stomatal density and aperture would change under Sµg, 2) these changes would differ between cultivars, and 3) these stomatal density changes in Sµg would correlate with pathogen ingression. We found that the four lettuce cultivars varied in stomatal density and aperture size under Sµg. Across the four cultivars, three had a significant increase in stomatal density on the adaxial surface while the fourth increased on the abaxial surface instead. Contrary to previous assumptions, *S. enterica* ingression was independent of stomatal density. Instead, cultivar selection appears to be a critical factor in selecting lettuce varieties for space habitation and missions, as red pigmented cultivars had less pathogen ingression compared to green cultivars. The other factor impacting ingression was microgravity, where ingression increased under Sµg. These findings enhance our understanding of how cultivar-specific genetics could be harnessed to improve food safety in both terrestrial and space-based crop cultivation.

## Results

### Simulated microgravity has significant impacts on stomatal density and ratio that vary between cultivars

To examine how Sµg affected seed leaf development and how cultivar specific genetic differences alter that response, we used fluorescent confocal microscopy and Image analysis to measure stomatal density and stomatal ratio between adaxial and abaxial stomata. Under normal gravity, lettuce microgreens had more stomata on the abaxial surface compared to the adaxial surface (p<0.01) (Figure 1A). SR analysis shows that when pooling all cultivars, SR increases under Sµg, with normal gravity mean SR at 0.85 and Sµg significantly increased to 1.18 (p<0.0001) (Figure 1B). However, examining individual cultivars shows variations in SR changes under Sµg (Figure 1C). The stomatal ratio in cultivars such as ‘Black seeded Simpson’ (BSS) and ‘Outredgeous’ (OUT) was similar under normal gravity and showed a similar increase in stomatal ratio when grown under Sµg. Cultivars such as ‘Flandria’ (FLA) and ‘Mascara’ (MAS) had a similar stomatal ratio when grown in normal gravity, however only MAS had a significant increase in stomatal ratio under Sµg. Overall, lettuce grown under Sµg had a significant increase in stomatal ratio, but the specific cultivars responded differently from each other.

**Figure 1:**
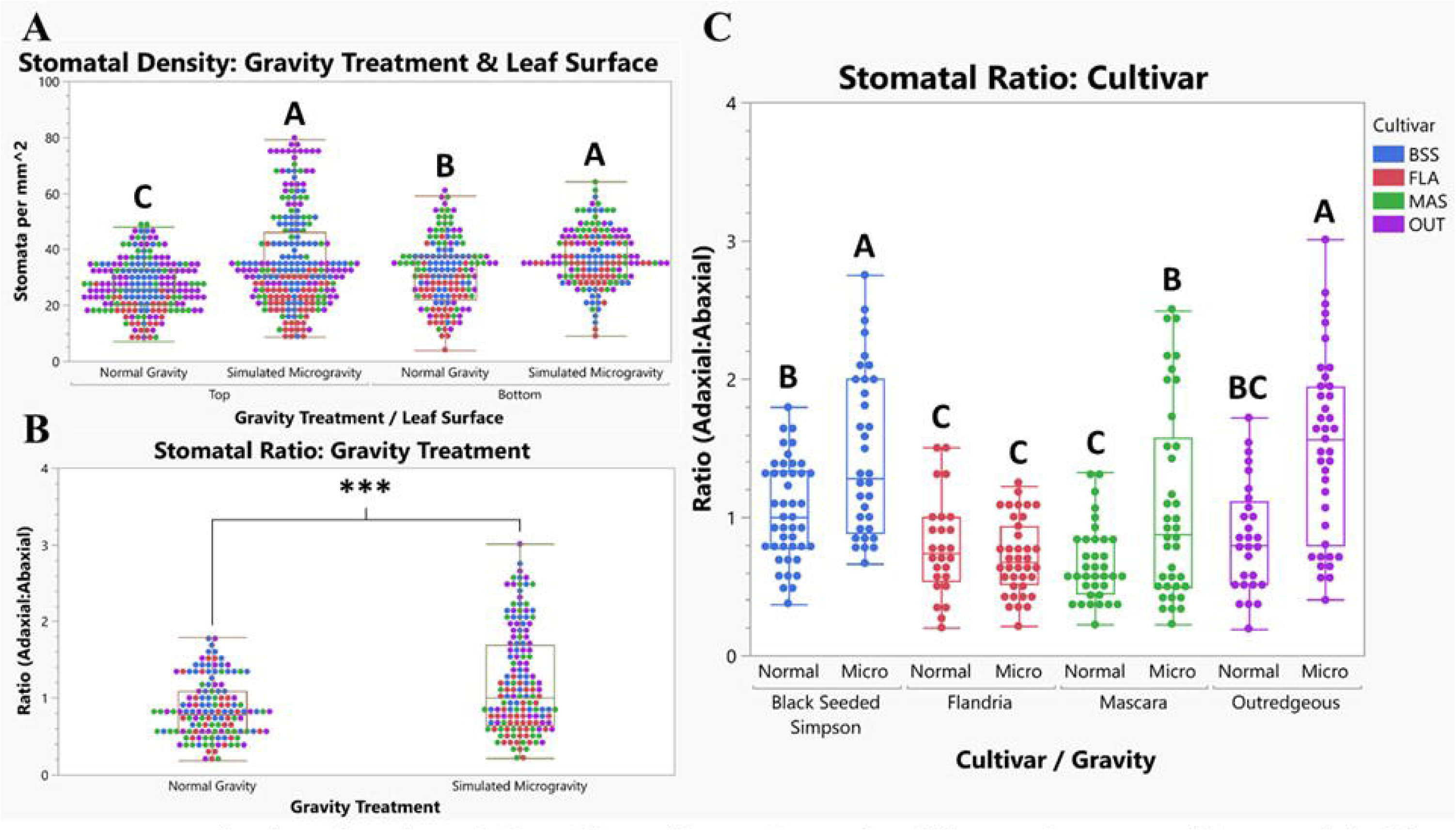
Stomatal ratio analyses in cotyledons of four cultivars of seven-day old lettuce microgreens. (A) Stomatal densities on the adaxial (top) and abaxial (bottom) surfaces under normal and simulated microgravity. (B) Stomatal ratios under normal and simulated microgravity. (C) Stomatal ratios are separated by cultivar and gravity treatment. *** indicate p<0.001. Connecting letters report generated by Student’s t tests for each graph where different letters indicate significance at 95% confidence.

Stomatal density on average was different between the four cultivars of lettuce (Figure 2). Stomatal density in lettuce grown with and without clinorotation at 4 RPM were significantly different between Sµg treatments, and each cultivar increased stomatal density under Sµg (p<0.05), though the surface where this change occurred was cultivar specific (Figure 2). On the adaxial surface of the cotyledon (Figure 2A) under normal gravity, three out of four cultivars started at approximately the same stomatal density (approximately 28 sq. mm^-1^). FLA grown at 0 RPM had significantly less stomata than the other cultivars on both sides of the cotyledon. On the adaxial surface, BSS and MAS had a similar significant increase in stomatal density in response to clinorotation. OUT had the strongest response to Sµg, with an average of 44 sq. mm^-1^ on the adaxial surface. Stomatal density did not increase in FLA (p=0.1505) under clinorotation on the adaxial surface. On the abaxial surface of the cotyledon, a very different response was noted (Figure 2B). The cultivars that had significant increases in stomatal density due to Sµg on the adaxial surface did not have similar increases on the abaxial surface. However, there was a significant increase in stomatal density due to Sµg in FLA. These differences between gravity treatments on the adaxial and abaxial sides are represented qualitatively via imaging in SOM Figure 1.

**Figure 2:**
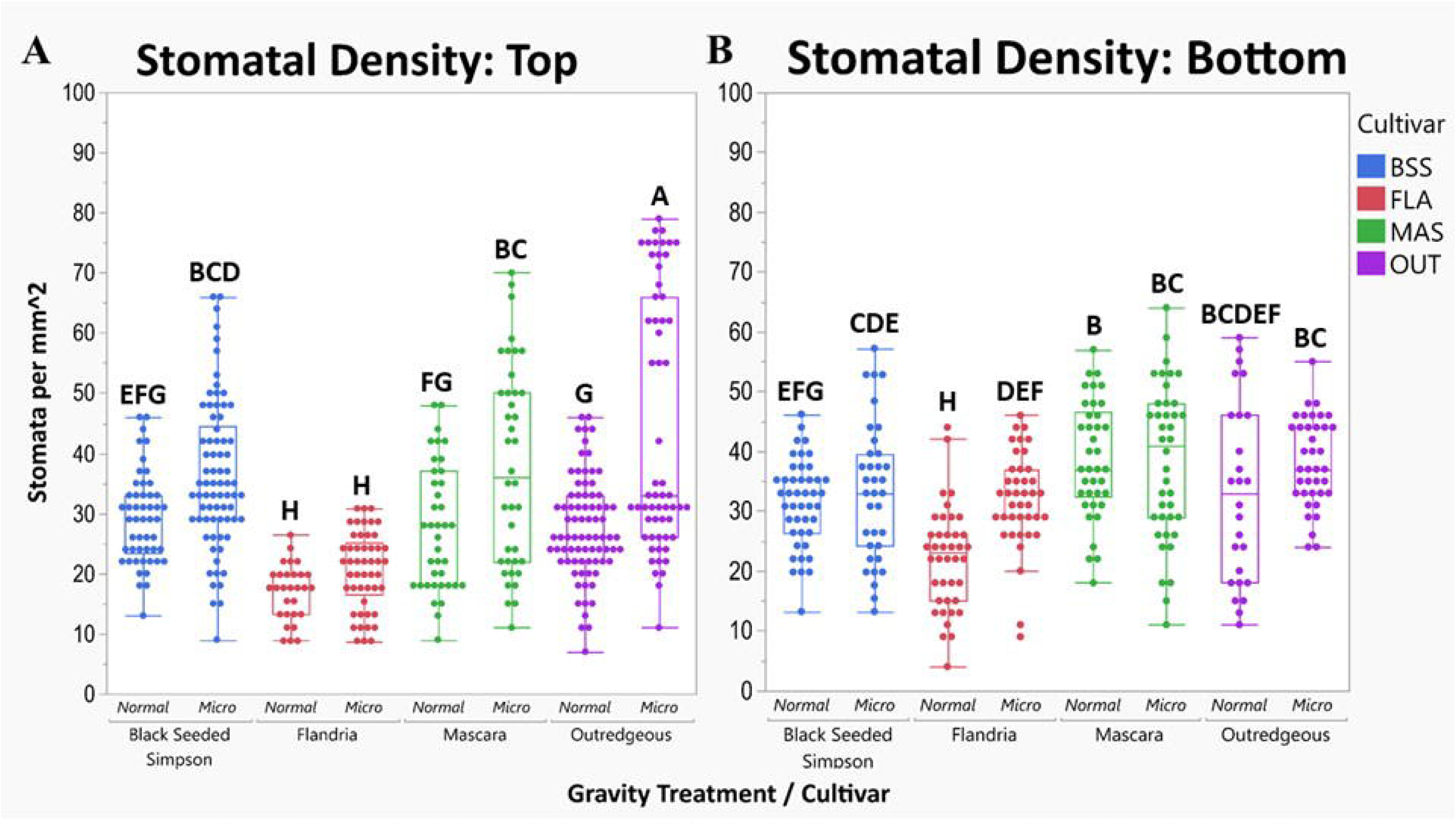
Stomatal densities on the (A) adaxial (top) and (B) abaxial (bottom) surfaces of cotyledons in four cultivars of seven-day old lettuce microgreens. Treatment codes along the x-axis indicate cultivar (BSS=’Black Seeded Simpson’, FLA=’Flandria’, MAS=’Mascara’, OUT=’Outredgeous’), Leaf surface examined (t=adaxial, b=abaxial), gravity treatment (0=normal gravity, 4=simulated microgravity through 2-D clinorotation at 4 RPM), and *S. enterica* application (- = sterile PBS mock application). Connecting letters generated between both graphs to compare stomatal densities between leaf sides by Student’s t tests where different letters indicate significance at 95% confidence.

### Simulated microgravity significantly alters stomatal aperture and defense between cultivars

Stomatal aperture was measured to see how Sµg affects stomatal defense against *Salmonella enterica* serovar Typhimurium 14028 or a mock treatment on the abaxial side. We also examined systemic innate stomatal responses to *S. enterica* inoculation by examining the adaxial surface of the cotyledon as well. All measurements were done at 9 hours post inoculation (hpi). There were significant differences in stomatal aperture and stomatal defense between cultivars, with OUT having on average the largest stomatal apertures, followed by BSS, FLA, and MAS (p<0.0001) (Figure 3). Stomatal aperture in lettuce grown with and without clinorotation at 4 RPM were significantly different between gravity and *S. enterica* treatments (F<0.0001), and different between cultivars (F<0.0001). BSS adaxial stomata under normal gravity did not change with the addition of pathogen to the abaxial surface; however, under clinorotation, stomata started more closed and opened with the pathogen inoculation (Figure 3A). The abaxial stomata under normal gravity closed with pathogen inoculation; however, under clinorotation, the stomata were more open without pathogen and opened further with the pathogen inoculation. In FLA on the adaxial surface, a closure response was noted under Sµg, but stomata opened with the pathogen challenge under normal gravity (Figure 3B). On the abaxial surface, no significant closure was observed regardless of pathogen exposure or exposure to Sµg. In MAS under normal gravity, adaxial stomata closed in response to pathogen exposure on the adaxial surface (Figure 3C). Under Sµg, stomatal aperture was lower than normal gravity controls and opened upon pathogen exposure, just like in BSS. However, all four adaxial treatments had a smaller stomatal aperture in MAS compared to BSS. On the abaxial surface, there was no significant closure with pathogen challenge regardless of the gravity treatment. In fact, stomatal aperture was similar to those seen in FLA. In OUT on the adaxial surface, a strong closure response was observed in both gravity treatments, though stomata remained more open under Sµg with pathogen treatment (Figure 3D). Adaxial stomatal aperture was notably larger under Sµg and no pathogen compared to normal gravity controls. On the abaxial surface, no closure response was observed regardless of the gravity treatment, though SA was smaller under Sµg. These responses to the four treatments on the adaxial and abaxial surfaces are qualitatively measured using imaging as shown in SOM Figure 2.

**Figure 3:**
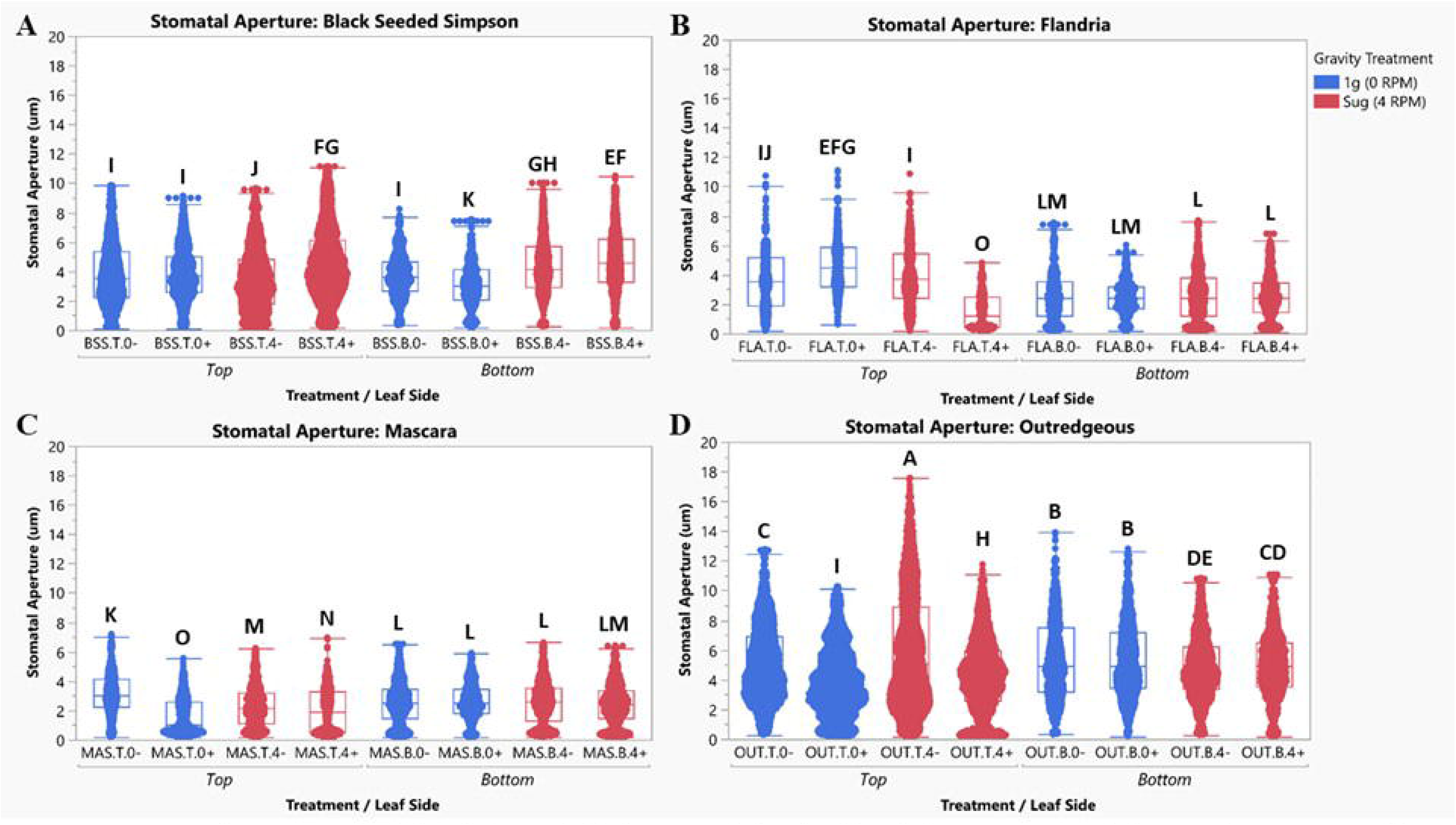
Stomatal Apertures in (A) ‘Black Seeded Simpson’, (B) ‘Flandria’, (C) ‘Mascara’, or (D) ‘Outredgeous seven-day old lettuce microgreen cotyledons. Treatment codes along the x-axis indicate cultivar (BSS=’Black Seeded Simpson’, FLA=’Flandria’, MAS=’Mascara’, OUT=’Outredgeous’), Leaf surface examined (t=adaxial, b=abaxial), gravity treatment (0=normal gravity, 4=simulated microgravity through 2-D clinorotation at 4 RPM), and *Salmonella enterica* application (- = sterile mock application + = *S. enterica)*. Connecting letters generated between all four graphs to compare cultivars by Student’s t tests, where different letters indicate significance at 95% confidence.

### Colony forming units (CFU) enumeration reveals cultivar- and pigment-specific changes to microbial population under simulated microgravity

To see how the stomatal phenotypic differences affected *S. enterica* populations, we performed CFU enumerations to look at both total and ingressed microbial populations per gram of fresh cotyledon tissue after 9 and 24 hpi. Each cultivar had different responses to Sµg (Figure 4). In BSS after 9 hpi, there were no significant changes to total or ingressed CFUs; however, after 24 hpi, both populations increased under Sµg (Figure 4A). Total and ingressed microbial populations increased after 15 hours of growth. In FLA after 9 hpi, there was a significant increase in total CFU but not the ingressed populations (Figure 4B). However, after 24 hpi there was no significant difference in total versus ingressed population. After 15 hours of growth, total microbial populations increase in both gravity treatments, however ingressed populations had no growth, regardless of the gravity treatment. In MAS after 9 hpi, there were no significant changes to total or ingressed CFUs; however, after 24 hpi, there was a significant decrease in the Sµg treatment due to a significant increase in microbial growth under normal gravity after 15 hours (Figure 4C). Ingressed populations remain constant regardless of incubation time or gravity treatment, similar to FLA. In OUT after 9 hpi and 24 hpi, there were significant decreases in total microbial populations under Sµg (Figure 4D). Under Sµg at 9 hpi, there were no significant differences between the total and ingressed populations, suggesting that most of the bacteria present were inside the apoplast. Under normal gravity, 15 hours of growth saw a significant decrease in ingressed microbial populations, which did not happen under Sµg. However, there was a significant increase in ingression between the normal and Sµg treatments at 24 hpi.

**Figure 4:**
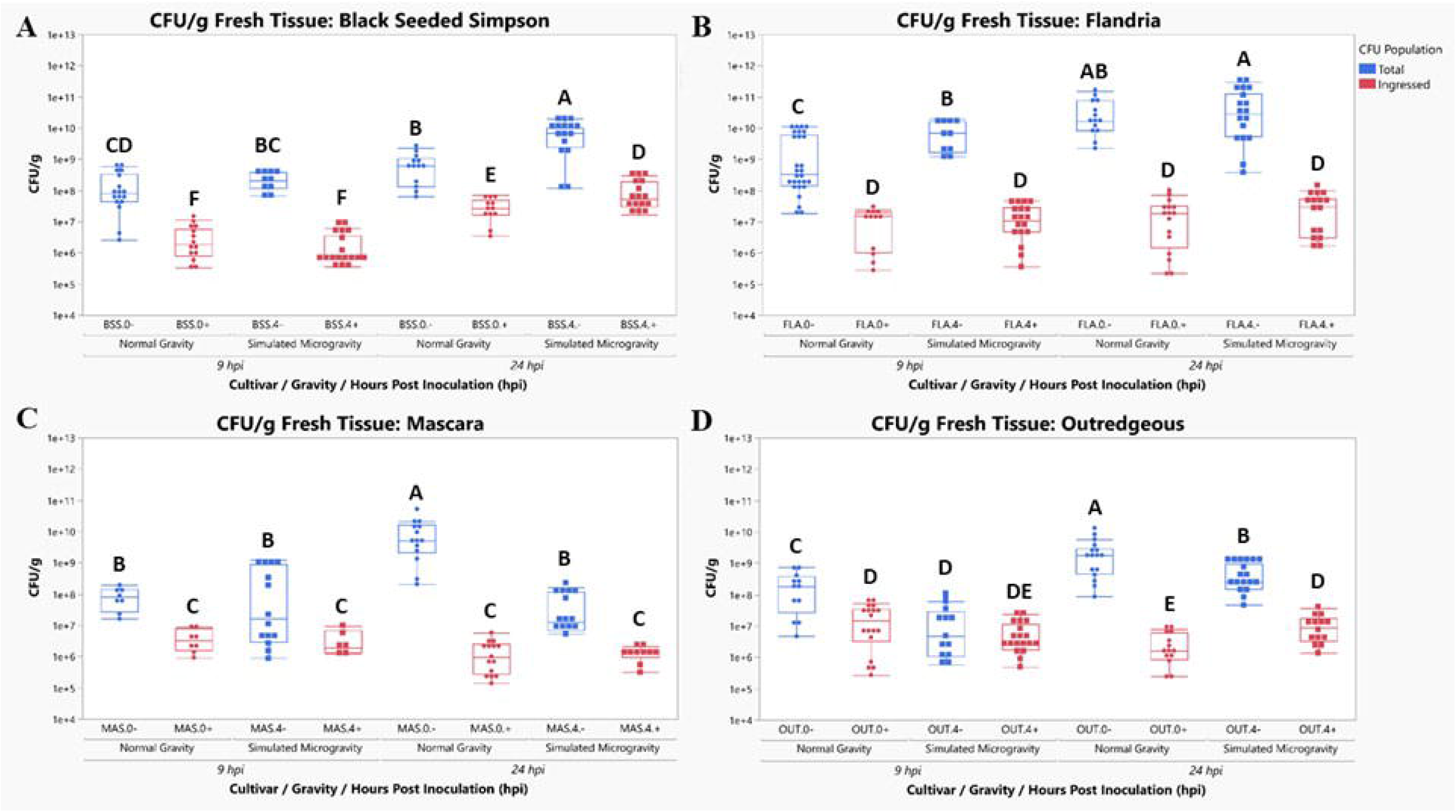
CFU per gram of fresh cotyledon tissue in (A) ‘Black Seeded Simpson’, (B) ‘Flandria’, (C) ‘Mascara’, or (D) ‘Outredgeous seven-day old lettuce microgreens. Treatment codes along the x-axis indicate cultivar (BSS=’Black Seeded Simpson’, FLA=’Flandria’, MAS=’Mascara’, OUT=’Outredgeous’) and surface sanitization condition (- = no surface sterilization, + = surface sterilization with 70% EtOH). Connecting letters generated for each graph by Student’s t tests, where different letters indicate significance at 95% confidence.

Removing cultivar from the equation and comparing normal gravity and Sµg grown plants, additional insights into how food safety may be impacted by spaceflight were observed. Total CFUs 24 hpi decrease under Sµg (p<0.05) (Figure 5A); however, ingression increases under Sµg (p<0.001) (Figure 5B), suggesting that more pathogens could be safe from standard food safety sanitization protocols in space. FLA and BSS are both cultivars that are primarily green in color, while MAS and OUT are cultivars that produce a strong red to purple pigmentation on their leaves. Separating the cultivars into ‘red’ and ‘green’ groups reveals that the two groups’ microbial populations respond differently to Sµg. Under normal gravity conditions, total pathogen populations did not differ significantly between the two groups. In contrast, under Sµg conditions, green cultivars exhibited a significant increase in pathogen population, whereas red cultivars showed a reduction (Figure 5C). Ingressed populations, however, are different between the two groups under normal gravity and Sµg, with red populations having significantly less compared to green (Figure 5D). However, both groups saw a significant increase in ingression under Sµg, with red cultivars having fewer ingressed CFUs when compared to the green pigmented cultivars.

**Figure 5:**
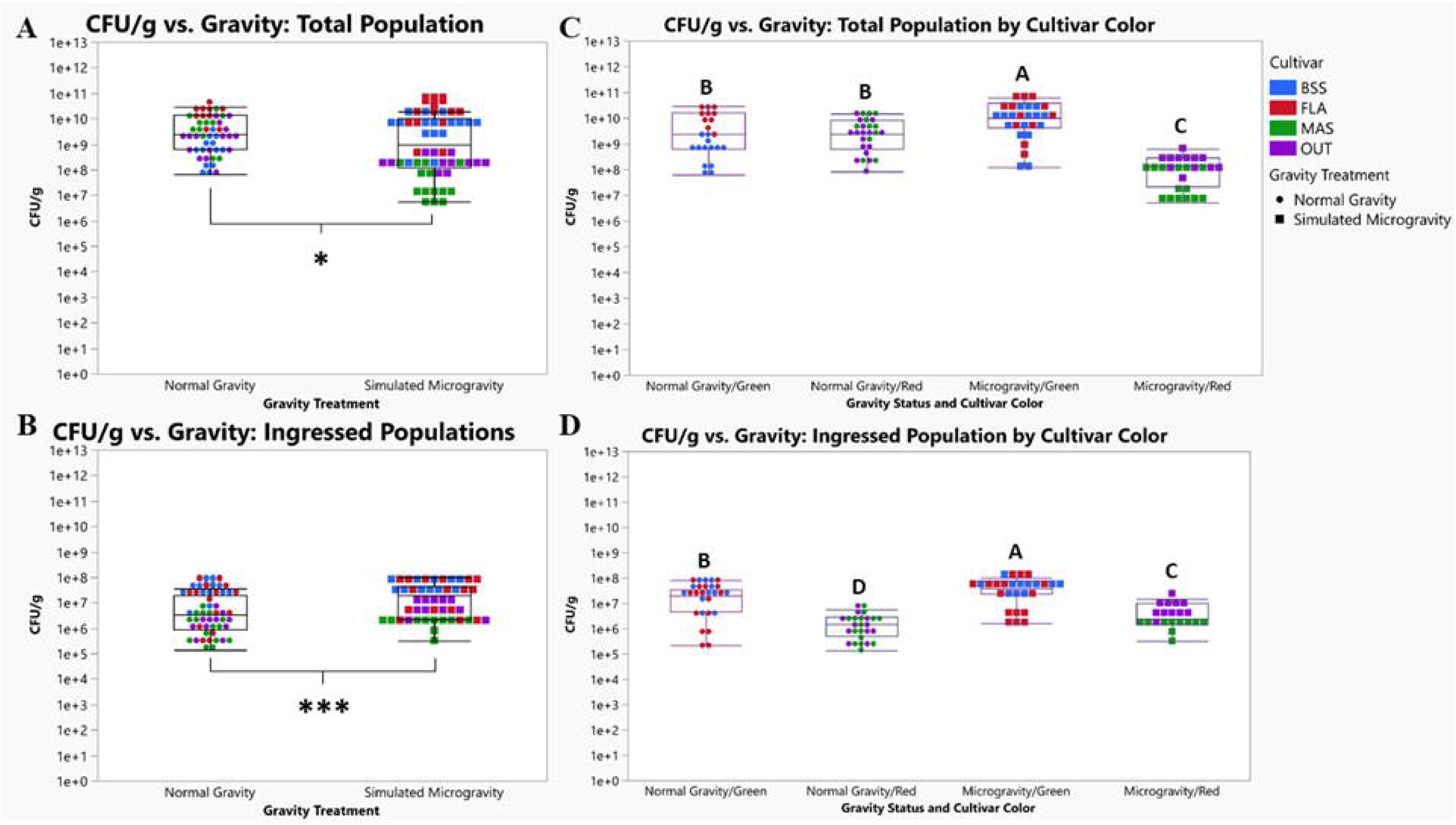
Analysis of CFU per gram of fresh cotyledon tissue by gravity treatment and cultivar color in total and ingressed microbial populations at 24 hpi. (A) All four cultivars comparing gravity treatments for total microbial populations. (B) All four cultivars comparing gravity treatments for ingressed microbial populations. (C) Cultivars are separated into red and green groups to compare gravity and color’s effects on total microbial populations. (D) Cultivars are separated into red and green groups to compare gravity and color’s effects on ingressed microbial populations. Asterisks denote significance where *= p<0.05 and ***= p<0.001. Connecting letters generated for each graph by Student’s t tests, where different letters indicate significance at 95% confidence.

### Stomatal density or aperture does not correlate with ingression, but density correlates with total microbial populations under simulated microgravity but not normal gravity

In a multivariate linear regression model examining the effects of cultivar and stomatal density on total and ingressed *S. enterica* CFUs under normal gravity and Sµg, cultivar is a significant predictor in all four models (Figure 6). However, stomatal density only improves the fit of the model of total microbial populations in plants grown under Sµg (Figure 6B). Plants grown under normal gravity (Figure 6 A&C) have a weaker fit for CFU populations compared to Sµg grown plants (Figure 6 B&D). Cultivar selection has a moderate impact on total (R^2^=0.53) and a weaker impact on ingressed populations (R^2^=0.44) in lettuce grown under normal gravity. Cultivar selection has a strong impact on total (R^2^=0.75) and ingressed (R^2^=0.71) populations in lettuce grown under Sµg. The fit of the total CFU under Sµg model increases significantly when stomatal density is factored into it (R^2^=0.81), leading to a strong positive correlation between stomatal density and total CFU. The impact of stomatal aperture on the total CFU was insignificant in all four models generated (data not shown).

**Figure 6:**
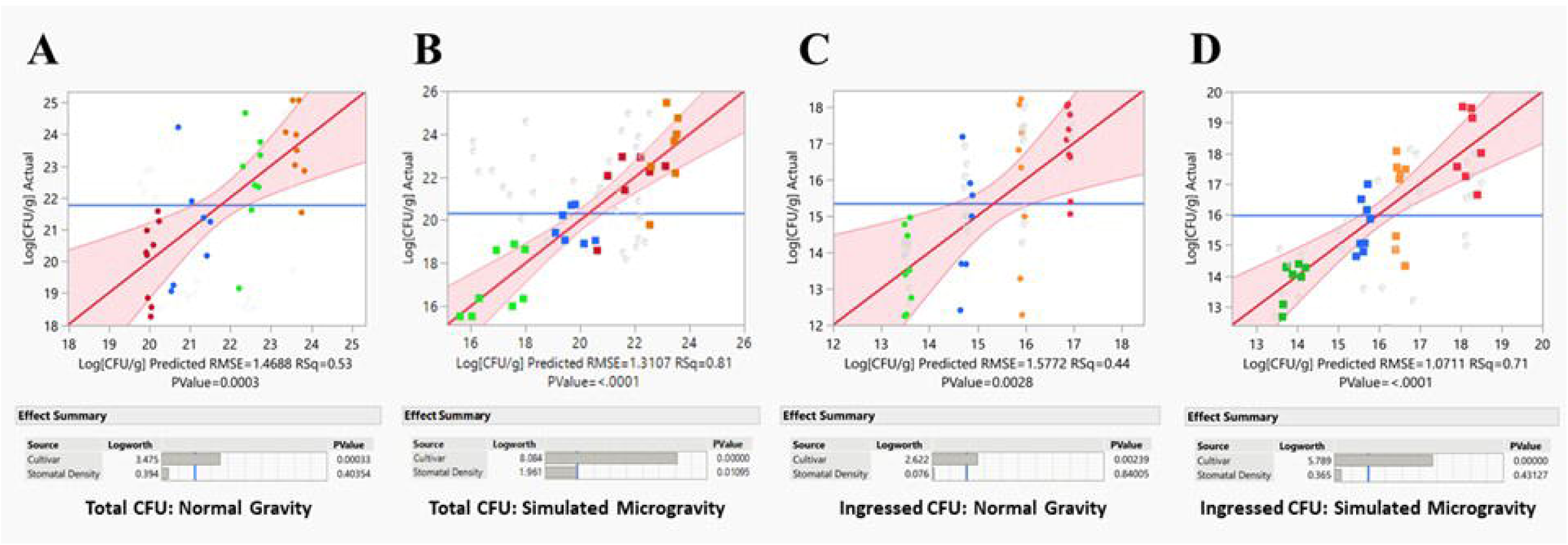
Multivariate linear regression models and effects tests for covariants cultivar and stomatal density on total and ingressed microbial populations under normal and simulated microgravity in four cultivars of seven-day old lettuce microgreens. (A) Total microbial populations under normal gravity. (B) Total microbial populations under simulated microgravity. (C) Ingressed microbial populations under normal gravity. (D) Ingressed microbial populations under simulated microgravity.

## Discussion

Here, we show the following key findings: stomatal density increased in four lettuce cultivars grown under Sµg, although the cotyledon surface on which this increase occurred was cultivar specific. Stomatal aperture and defense against *S. enterica* were also affected by Sµg in a cultivar-dependent manner. Red-pigmented cultivars ‘Mascara’ and ‘Outredgeous’ exhibited decreased total microbial persistence under Sµg, whereas the green cultivars ‘Black Seeded Simpson’ and ‘Flandria’ showed increased microbial populations; however, both groups experienced significantly greater pathogen ingression under Sµg conditions. Notably, stomatal density and stomatal aperture did not significantly influence ingression, whereas cultivar selection and Sµg exposure did. Nonetheless, stomatal density improved predictions of total microbial population under Sµg. Taken together, these results suggest that simulated microgravity conditions encountered during spaceflight can significantly alter stomatal development and function, and that cultivar selection has significant implications for plant–microbe interactions and food safety in lettuce production systems on, and possibly off, Earth.

The present work aligns with previously published research demonstrating that lettuce cultivar BSS grown under Sµg using 2-D clinorotation at 4 RPM exhibit significant alterations in stomatal aperture and *S. enterica* ingression (Totsline et al., 2024). However, these results may contradict others. A study by Monje et al. (2005) using wheat suggests that true microgravity itself may not substantially alter stomatal function, though they did not examine stomatal aperture. In that work, plants grown on Earth under simulated space station environmental conditions but maintained under normal gravity showed no significant differences in other stomatal function metrics, evapotranspiration rates or net photosynthesis, compared to plants cultivated aboard the Mir Space Station. Further evidence that Sµg can influence stomatal behavior comes from a study on the C_3_ monocot *Chlorophytum comosum*, where plants grown in a random positioning machine exhibited altered stomatal function. Specifically, dark-adapted plants under Sµg conditions maintained open stomata, in contrast to closed stomata in dark-treated controls grown under normal gravity (Treesubsuntorn et al., 2020). Stomatal apertures were not measured during day or night cycles in this study. These findings indicate that Sµg can significantly impact stomatal function, as was shown in our study. The current body of literature in this area remains limited, emphasizing both the novelty of the present study and the need for continued research.

The prior work examining how stomatal density changes under spaceflight or Sµg conditions is also extremely limited and contradicts this study. In *Prunus jamasakura*, plants grown with and without 3-D clinorotation at 1 RPM for four weeks showed no significant changes in stomatal density on the abaxial leaf surface (Sugano et al., 2002). *P. jamasakura* is likely fully hypostomatic; however, it remains unclear whether growth under microgravity conditions can modify stomatal distribution on true or cotyledonary leaf surfaces to an extent that induces a transition from hypostomatic to amphistomatic leaf morphology. There is some evidence that Sµg can influence stomatal absence/presence in non-leaf tissues, where Sµg at 20 RPM has been shown to eliminate stomata from the stems of dark-grown tomato seedlings (Terna et al., 2017). This finding supports the rationale for examining both leaf surfaces in microgravity experiments, even in species that are typically hypostomatous under terrestrial conditions. Consistent with this reasoning, the present study did not detect significant changes in abaxial stomatal density in three out of four lettuce cultivars examined. In contrast, significant increases in stomatal density were observed on the adaxial cotyledon surface in those same cultivars under Sµg. Stomatal density has been shown in *Arabidopsis thaliana* to be semi-independently regulated between the adaxial and abaxial leaf surfaces (Jalakas et al., 2024), a pattern that is consistent with the findings of the present study. Although stomatal density increased across all four lettuce cultivars under Sµg, the specific leaf surface on which this increase occurred was cultivar dependent. Jalakas et al. (2024) investigated this differential regulation using abscisic acid (ABA) synthesis, catabolism, and sensing mutants, along with known stomatal development mutants in *A. thaliana*. Their results showed that key regulators of stomatal development—*SDD1, EPF1/EPF2,* and *TMM*—preferentially suppressed stomatal formation on the abaxial surface, whereas increased ABA concentration and *ERL2* expression suppressed development on the adaxial surface. The observed increase in adaxial stomatal density in three lettuce cultivars in the present study could therefore hypothetically be linked to reduced ABA levels under Sµg conditions.

Supporting this possibility, Carman et al. (2015) reported significantly lower ABA concentrations in wheat crowns and leaves grown in space compared to ground-grown controls. However, interpretation of these findings is complicated by substantial differences in growth conditions between spaceflight and ground controls, including variation in substrate, photoperiod, humidity, temperature, and light intensity, which may confound ABA-related effects. Similar limitations are present in other studies examining ABA dynamics under spaceflight conditions. Moreover, the role of ABA in regulating stomatal density remains unresolved, even under terrestrial conditions. While elevated ABA levels have been shown to negatively regulate stomatal density in *A. thaliana* (Jalakas et al., 2024; Wei et al., 2021), contrasting evidence indicates that ABA can positively regulate stomatal density in basil (Jensen et al., 2018) and *A. thaliana* (Lake and Woodward, 2008). These discrepancies highlight the complexity of ABA-mediated stomatal development and suggest that both species-specific responses and environmental context play critical roles. Overall, further research conducted under controlled terrestrial and spaceflight conditions will be necessary to establish clear trends and mechanistic understanding of ABA’s role in stomatal density regulation.

The absence of a correlation between stomata aperture and ingression wasn’t unexpected. Roy et al. (2013) found that high humidity environments, like the one created by growth in petri dishes, can impair closure against *S. enterica* but not *Escherichia coli* O157:H7. However, stomatal closure occurred on the adaxial surface where *S. enterica* was not applied, suggesting a systemic response to *S. enterica* that was not impaired by high humidity. The impaired stomatal defense response on the abaxial surface may be explained by the findings of Johnson et al. (2020), who demonstrated that *Salmonella enterica* serovar Typhimurium employs its Type III secretion system (T3SS), a virulence mechanism classically associated with animal host infection, to suppress stomatal defense responses in lettuce. *S. enterica* T3SS *hilD* and *sseB* mutants had a stronger stomatal defense response against the pathogen, in addition to reduced persistence inside the leaf. *S. enterica* has also been shown to synthesize auxin and induce auxin biosynthesis in *A. thaliana* to re-open stomata (Fochs et al., 2025), further highlighting why stomatal defense was unlikely to impair ingression against this particular pathogen.

The absence of a correlation between stomatal density and ingression, however, was unexpected, with cultivar selection being the strongest predictor of ingression CFU values instead. Cultivar selection impacting *S. enterica* proliferation has been observed in other lettuce studies (Erickson et al., 2019; Kim et al., 2018; Liu et al., 2023). Erickson et al. (2019) observed that cultivars rich in antimicrobial compounds, specifically phenols and other antioxidants, had a significant strong inverse relationship with pathogen internalization. Generally, red pigmented cultivars have more phenols and antioxidant capacity compared to green lettuce cultivars (Kim et al., 2018), which could explain in part why the red cultivars had less ingression. The presence of these compounds can impact surface microbial population as well. Liu et al. (2023) found that red ‘Mascara’ oak leaf lettuce had more flavonoids, a major class of phenols, and anthocyanins than green ‘Parris Island Cos’ romaine. The red cultivar showed significantly lower recovery of *S. enterica* serovars Newport and Typhimurium from produce rinses compared with the green cultivar. ‘Mascara’ was included in this study and had the lowest total and ingressed microbial populations, suggesting that it may also be the most nutrient dense cultivar of the four cultivars studied in the present work; however, this has not been confirmed. Why microgravity impacts total and ingressed microbial populations is unclear, and existing literature that could explain this difference is sparse. Microbial ingression could increase during microgravity due to impaired closure in darkness. While the current study did not examine stomatal closure in dark adapted stomata, impaired closure during the dark period has been observed in both true (Monje et al., 2000) and simulated microgravity (Treesubsuntorn et al., 2020). *S. enterica* have been observed to be attracted to open stomata; however, stomata forced open in lettuce during the dark period had no significant impact on ingression, suggesting that *S. enterica* cells are attracted to compounds produced by photosynthetically active cells (Kroupitski et al., 2009). More research is needed to understand why ingression is increasing under microgravity.

Changes in microbial populations under Sµg may be due to suppressed plant immune systems combined with pathogens having improved fitness in both simulated and true microgravity environments (Totsline et al., 2023). Recent analyses of lettuce grown in space have found no human pathogens persisting in and on the plants (Khodadad et al., 2020); however, it is quite possible that these pathogens could contaminate space environments in the future (Colorado et al., 2021), and proactive research in the area could reduce the severity of outbreaks should they happen. While research is ongoing in the field of food safety in space and existing research is limited, what exists suggests that infections with foodborne illnesses in space may be more severe. In *S. enterica* serovar Typhimurium grown at 25 RPM to create a Low Shear Modeled Microgravity (LSMMG) environment, mice exposed to the pathogen grown under Sµg had a 20% survival rate 10 days post infection compared to 60% from the normal gravity grown pathogen (Nickerson et al., 2000). In similar culture conditions, LSMMG *S. enterica* had improved adherence, invasion, and intracellular survival in a 3-D biomimetic model of human colonic epithelium containing macrophages (Barrila et al., 2022). To the authors’ knowledge, only the Totsline et al. (2024) study examines the foodborne pathogen–plant interaction in the context of microgravity. Thus, much is unknown about how any foodborne pathogens behave in microgravity before they reach a human host in the spaceflight environment.

It is important to note that the method used to simulate microgravity in microbes—that is, through the creation of a low shear environment in liquid cultures—is not how microgravity is simulated in plants. Any plant–microbe interaction experiment performed on Earth cannot simulate microgravity on both the microbe and the plant at the same time due to these mechanistic differences in microgravity simulation. LSMMG uses rotation to simulate fluid dynamics like those that microgravity create in liquid cultures. They have been shown to decrease the effects of Earth’s gravitational force (10^0^ x g) significantly on the culture fluid (10^-2^ x g), though not quite to true spaceflight levels (10^-4^ – 10^-6^ x g) (Higginson et al., 2016). Simulating microgravity in plants is primarily through preventing starchy statoliths from settling in the root tip, the predominant hypothesis surrounding gravity sensing and responses in plants (Kiss, 2000). This can be done with one-, two-, or three-axial clinostats, random positioning machines, magnetic levitation, and centrifuges (Kiss et al., 2019). Slow rotation (2–4 RPM) with a one-axial clinostat, like the one used in this study, was shown to correspond to effective randomization of statoliths position (Dedolph and Dipert, 1971) to simulate microgravity. However, in true microgravity, statoliths have been shown to cluster in proximal parts of the cell due to changes in actomyosin function in space (Muthert et al., 2020), suggesting that this method of simulating microgravity is not a perfect analog. In fact, every method to simulate microgravity on Earth, regardless of the organism studied, introduces artifacts; thus, conclusions drawn from experiments conducted in ground-based facilities (GBFs) must be verified in space. However, true microgravity experiments are expensive and competitive due to limited availability, so data collected in GBFs can provide valuable insights into what experiments are worth the resources to send into space.

## Conclusion

Foodborne illness remains a significant public health concern, particularly as fresh produce consumption continues to increase worldwide and in controlled environments such as the ISS. *Salmonella enterica* is a leading cause of hospitalization and mortality associated with foodborne pathogens in the United States (Thomas et al., 2024). The significance of the present work extends to both terrestrial and space-based agricultural systems (Figure 7). Notably, lettuce cultivars that develop red pigmentation exhibited reduced total microbial populations and lower populations of ingressed bacteria, which are less accessible to conventional sanitization methods. However, the underlying mechanisms driving these differences remain unclear. Importantly, Sµg increased pathogen ingression in both red and green lettuce cultivars, suggesting an elevated risk of foodborne illness during long-duration space missions if contamination of space-grown produce occurs. In addition to effects on plant–microbe interactions, Sµg also increased stomatal ratio and stomatal density in lettuce, which may influence broader plant stress responses in this novel and challenging environment. Despite these changes, variation in stomatal density and stomatal aperture was strongly cultivar-dependent and did not directly correlate with pathogen ingression. This indicates that stomatal traits alone are not primary determinants of *S. enterica* entry into cotyledon tissue under Sµg conditions. Consequently, plant breeding strategies aimed at improving food safety—whether on Earth or in space—should prioritize other physiological or biochemical traits rather than targeting stomatal density.

**Figure 7.**
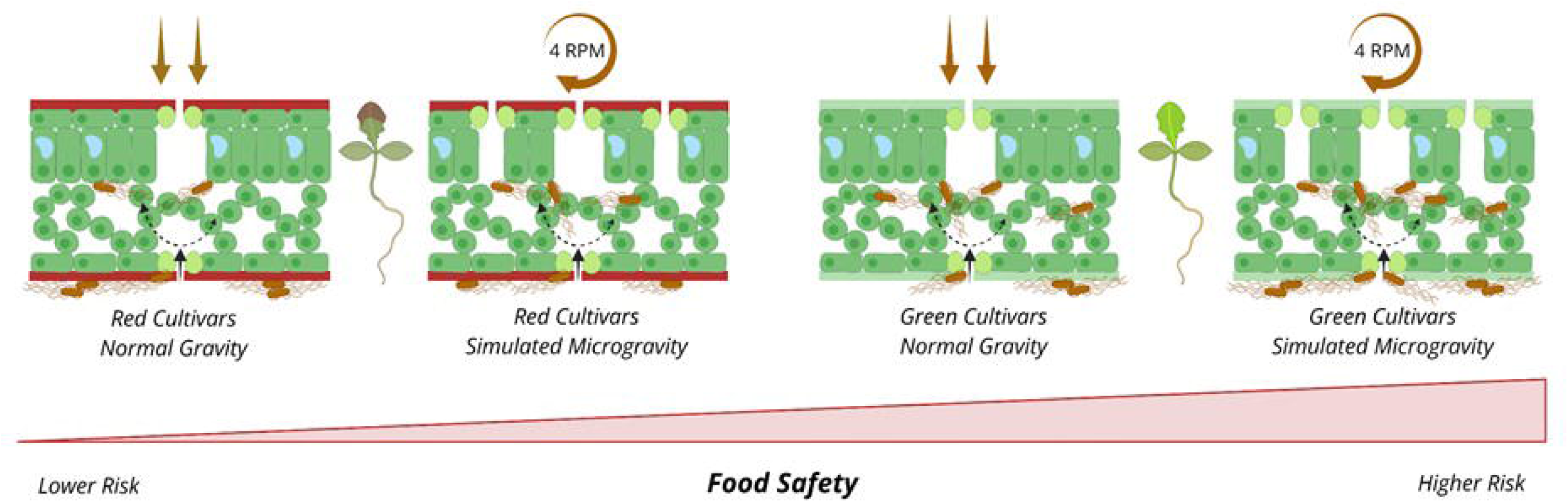
Conceptual model summarizing key findings linking stomatal development to food safety outcomes in lettuce microgreens. Simulated microgravity increases stomatal ratio independent of cultivar pigmentation and significantly alters both total surface and internalized (ingressed) populations of Salmonella enterica. Total microbial populations increase under simulated microgravity in green cultivars but decrease in red cultivars, whereas internalized populations increase across both types. Notably, red cultivars exhibit lower baseline levels of internalized pathogens and, despite microgravity-induced increases, maintain lower internalized populations than green cultivars under comparable conditions. The model supports the hypothesis that cultivar-dependent traits modulate food safety risk, with red cultivars representing a potentially lower-risk phenotype under both terrestrial and simulated microgravity conditions.

## Materials and Methods

### Plant materials and growing conditions

Seeds from four cultivars of lettuce, green leaf ‘Black Seeded Simpson’ (BSS), green butterleaf ‘Flandria’ (FLA), red leaf ‘Mascara’ (MAS), and red romaine ‘Outredgeous’ (OUT), were stored long term in the dark at 4 ℃ in their original seed packet. Seeds were surface sterilized with 9 mL of a 2:1 commercial bleach to DI water solution in a 15 mL conical tube, shaken for 9 minutes, the bleach solution was decanted, and the seeds were rinsed with 9 mL of sterile DI water 5 times. 40 mL of 0.8% Phytagel media supplemented with 1% w/v sucrose and 1⁄2 strength Murashige and Skoog plant growth media was poured in 90 x 90 cm square petri dishes and were allowed to solidify in an aseptic laminar flow hood. The top 30 mm of solidified media was removed using a sterilized flatula to allow additional space for vertical growth and to create a flat surface upon which seeds could germinate. Four seeds were placed approximately 20 mm apart using a sterilized flatula, and plates were sealed using micropore tape. The plates were positioned vertically within a two dimensional slow-rotating clinostat designed by Tania Wiest and synthesized by Alfred Lance of the University of Delaware machine shop (SOM Figure 3). Three rows of LEDs (KYUQV Amazon) with red: blue lights at a 5:1 ratio were in each growth unit and were programmed to a light intensity of 150 μmol m^−2^/s with a 12-h photoperiod. Microgreens were grown for 7 days after the first true leaf is consistently present either with continuous rotation at 4 RPM to simulate microgravity or without rotation to provide normal gravity. For each treatment three plates were sampled, with one cotyledon sampled from the three largest microgreens (n=9 cotyledons per treatment).

### 2.2 Bacterial culture

*Salmonella enterica* subsp. *enterica* serovar Typhimurium GFP 14028GFP^TM^ supplied by the American Type Culture Collection was stored in 40% glycerol in −80 °C freezer for long-term storage and cultured using protocols established by Totsline et al., (2024). Before use in experiments, bacteria were streaked from glycerol stocks onto LB (Miller Broth) with 0.8% Agar and 100 µg/mL ampicillin and incubated at 37 ℃ for 12 h. Following solid media incubation, a single colony forming unit (CFU) was transferred using a sterile inoculation loop in a 15 mL conical tube containing 5 mL LB (Miller Broth) with 100 µg/mL ampicillin. Liquid cultures were incubated in an orbital shaker set to 250 RPM at 37 °C for 12 h. Following liquid media incubation, 2 mL of culture was transferred to a microcentrifuge tube, then a bacterial pellet was obtained by centrifugation at 4000 x g for 5 min. The pellet was resuspended by vortexing and washed twice with 2 mL of phosphate buffered saline (PBS) at a pH of 7.4, followed by a final suspension in PBS. The optical density was measured at 600 nm (OD_600_) with a Bio-Rad SmartSpec + spectrophotometer (Bio-Rad Inc.) and adjusted to an OD_600_ of 0.8 with a final concentration of 10^8^ cells/mL. All experiments involving *S. enterica* Typhimurium used the methods described above and experiments were immediately conducted following final adjustment of OD_600_.

### 2.3 Foliar application of Salmonella enterica

Following final adjustments of OD_600_, a 10µL sized inoculation loop was flame sterilized, cooled, then submerged in either the suspension of *S. enterica* or sterile PBS for mock inoculations. Care was taken to ensure the loop was full of inoculum to ensure consistent application of culture between cotyledons. The abaxial surface of both cotyledons of the largest three microgreens in a petri dish were gently rubbed with the inoculation loop. The inoculation loop was flame sterilized between applications. Plants were then returned to the clinostat (SOM Figure 1) either in rotating or stationary growth units depending on the treatment, for either 9 hours for stomatal analyses and CFU enumeration or 24 hour for another CFU enumeration.

### 2.4 Stomatal density and aperture analysis

9 hours following inoculation with either *S. enterica* or sterile PBS, the petri dishes were removed from the clinostat and a single cotyledon was sampled from each of the three inoculated plants from the three dishes for each treatment (n=9 cotyledons per treatment). Cotyledons were immediately excised from petioles using sterile forceps and submerged in 500 µL fixative solution (2% Glutaraldehyde + 2% paraformaldehyde + .05% Triton X-100 in PBS) in a 24-well plate. The plate was transferred to a vacuum desiccator and a vacuum of -0.1 MPa was generated for 20 seconds, then the vacuum was released and the chamber returned to ambient pressure. This step was repeated twice, the third time the vacuum was maintained for an hour. The cotyledons were then allowed to sit in the solution overnight at room temperature in the dark. The fixative solution was decanted, then 500 µL of 0.1 mg/mL solution of propidium iodide was added to each well and cotyledons were stained in the dark for 9 minutes. Then the propidium iodide solution was decanted and 500 µL of PBS was added to rinse the cotyledons. The PBS was decanted and the rinsing step was repeated. The final rinse was not decanted to improve ease of transferring samples for mounting. Cotyledons were mounted either adaxial or abaxial side down in separate glass bottomed Nunc chambers with 100 µL of 80% glycerol and glass blocks placed on top.

Cotyledons were imaged with inverted-confocal laser microscopy using a Leica STELLARIS 8 tauSTED located in the University of Delaware Bioimaging Core. Five random images were taken of each cotyledon surface to obtain images for stomatal density and stomatal aperture analysis (n=5 images for stomatal density; n=4-45 stomata per image to measure for stomatal aperture, variation dependent on treatment). Images were captured using a 20x glycerol immersion magnification objective + 1.28x digital zoom and a numerical aperture of 1.20. Images were acquired following a protocol established by Totsline et al., (2024) with 2176 × 2176 pixels per frame, pixel size 0.049 µm, bit-depth 12, Airy Units set to 1.0, a unidirectional X-axis speed of 400 Hz, pixel dwell time of 1.35 µs, frame rate of 0.331/s. and a physical length of 106.02 × 106.02 μm per frame. Propidium iodide stain was excited with a 489 nm laser at 2% intensity through a 488/561 bandpass filter with the emission spectra set to 509–562 nm and digital gain set to 20%. The GFPmut3 protein was excited with a 504 nm laser at 7% intensity through a 360/40 bandpass filter with emission spectra set to 588–725 nm and digital gain set to 2.5% (Totsline et al., 2024). Following imaging, stomatal apertures and densities were measured using ImageJ. The ellipse tool was used to trace the stomatal aperture, and the minor axis was calculated to provide a precise measurement of the width of the stomatal pore, or aperture. Each stoma in each image was measured to prevent bias and to obtain counts for stomatal density (sq. mm^-1^) and stomatal ratio (ratio of adaxial and abaxial stomatal densities).

### 2.5 CFU Enumeration

A single cotyledon was sampled from the three largest inoculated plants in 3 square petri dishes (n=9) at 9- and 24-hpi with *S. enterica*. Cotyledons were excised from petioles with sterile forceps, put into weighed microcentrifuge tubes, and weighed again to obtain fresh mass. Some treatments involved a surface sterilization step to examine only ingressed populations of *S. enterica* following a protocol by Jacob et al., (2017). For surface sterilization, 1000 µL of 70% ethanol was added to each tube, capped, shaken, and incubated at room temperature for 2 minutes. Next, the cotyledons were removed from the ethanol solution and transferred to tubes filled with 1000 µL of sterile PBS, capped, shaken, and incubated at room temperature for 1 minute. Afterwards, surface sterilized tissues were treated the same as the non-sterilized tissues.

Cotyledons were then transferred to a microcentrifuge tube with 100 µL sterile PBS and a sterile metal ball bearing. Samples were then transferred to a TissueLyser II (QIAGEN, Germany) for homogenization at 30 Hz for 30 seconds. Samples were redistributed in the homogenizer, then homogenized at 30 Hz for 30 seconds again. The homogenized samples were serially diluted at a ratio of 1:10 by removing 10 µL of solution and adding it to a microcentrifuge tube containing 90 µL of sterile PBS for a 10^-1^ dilution. This was repeated to reach a 10^-5^ dilution. Each dilution tube was mixed via pipetting five times and two replicates of 10 µL was removed and drop plated in one square of a 90 x 90 grid petri dish of LB (Miller Broth) with 0.8% agar and 100 μg/mL ampicillin. Petri dishes were sealed with micropore tape and incubated for 12 h at 37 °C. Single colony forming units (CFUs) were counted and CFUs per gram of fresh cotyledon tissue were calculated

### 2.6 Statistical Analysis

Data was analyzed using a one-way ANOVA to determine the effect of treatments, and a student’s t multiple comparisons test was used to compare means of stomatal aperture, stomatal density, stomatal ratio, and CFU enumerations by using JMP software (JMP v.14; SAS Institute Inc., Cary, NC) at a significance level of p < 0.05. JMP software was also used to create multivariate linear regression models. Since stomatal density and aperture data sets were larger than the CFU data sets, to assess the correlations between stomatal densities or apertures to CFU populations, average densities/apertures for each leaf replicate (n=9) were aligned with a random CFU value (n=9) for each treatment to generate a scatterplot utilized for multivariate linear regressions.

## Supporting information

SOM Figure 1

SOM Figure 2

SOM Figure 3

## Acknowledgements

TAMW is grateful to the current and past members of the Bais lab for their support and ideas throughout this project. HPB acknowledges NASA-EPSCoR (DE-80NSSC20M0253) for funding support.

## Author Contributions

TAMW and HPB designed the experiments, TAMW performed all experiments, TAMW analyzed the results, TAMW and HPB wrote the first draft of the manuscript. All authors have read and agreed to the published version of the manuscript.

## Legends to Supplementary Online Figures

**SOM Figure 1:** 2-D Clinostat designs (A-C) and built machine (D) utilized for this study. (A) Side view of clinostat, showing the two rotating and two stationary growth units, and the desired sizes for each one. Below that shows orientation of three strings of lights used on both sides of the interiors of the growth units. (B) Front view of clinostat. (C) Schematic for growth units, designed for versatility with three kinds of experimental designs. The one utilized for this study was to use four square 10x10-petri dishes on both sides of the glass adjusted to sit in the middle of the growth unit (eight total dishes). The off-center support screw also allows for a set up with two 10x10 petri dishes on the shorter right side and two 12x12 petri dishes on the left side to allow for older *in vitro* plant studies. With a similar set-up on the other side of the glass, this would also allow eight plates at a time as well. The width of these growth units and the removal of the glass in the middle would allow for 10x10 cm plant pillows or specialized pots to be affixed to the bottom of the growth units, allowing for larger plants grown in a greenhouse setting compared to *in vitro.* This arrangement would support four pots per growth unit. (D) Actual clinostat built with these schematics used in this study.

**SOM Figure 2:** Fluorescent confocal images at 200x magnification of average stomatal densities on adaxial and abaxial surfaces of cotyledons in four cultivars of seven-day old lettuce microgreens with (4 RPM) and without (0 RPM) simulated microgravity. Leaves were imaged with inverted-confocal laser microscopy using a Leica STELLARIS 8 tauSTED located in the University of Delaware Bioimaging Core and analyzed using ImageJ.

**SOM Figure 3:** Fluorescent confocal images at 200x magnification of average stomatal apertures on adaxial and abaxial surfaces of cotyledons in four cultivars of seven-day old lettuce microgreens with (4 RPM) and without (0 RPM) simulated microgravity and with (+) or without (-) inoculation with GFP-tagged *Salmonella enterica.* Leaves were imaged with inverted-confocal laser microscopy using a Leica STELLARIS 8 tauSTED located in the University of Delaware Bioimaging Core and analyzed using ImageJ.

