## Supplementary figures and images for "Simulated Microgravity Induces Cultivar-Specific Changes Affecting *Salmonella enterica* Ingression Independent of Stomatal Physiology"

### SOM Figure 1

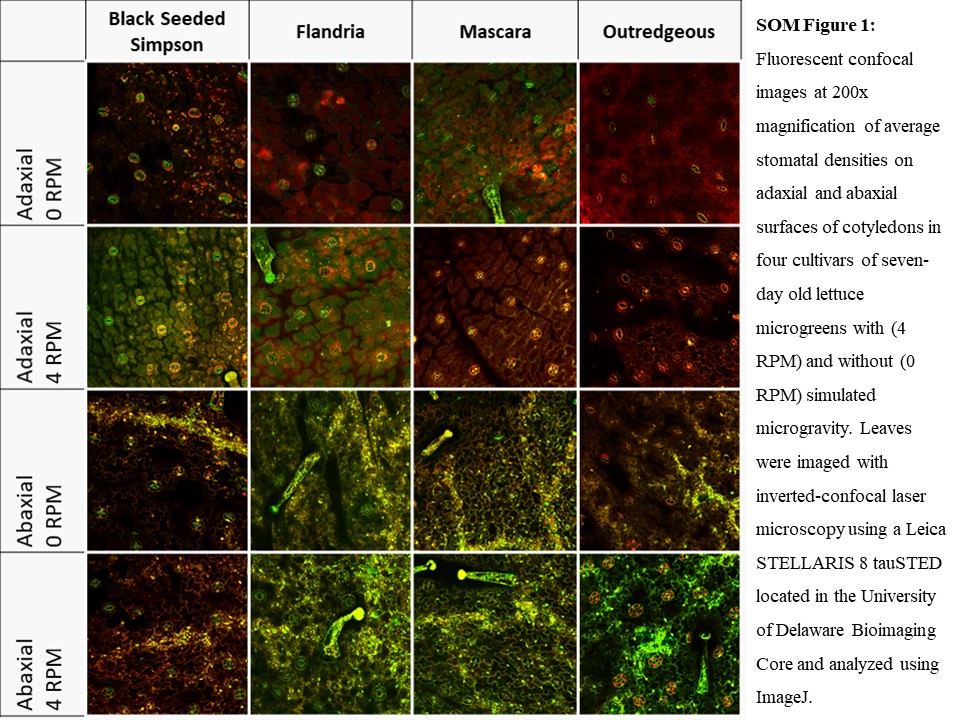

### SOM Figure 2

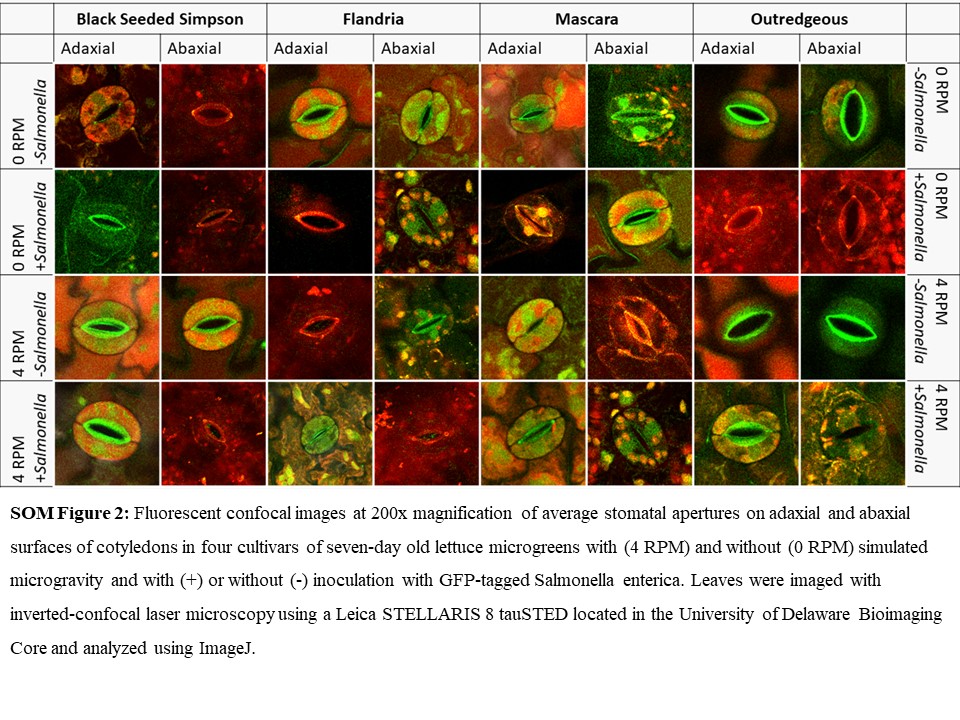

### SOM Figure 3

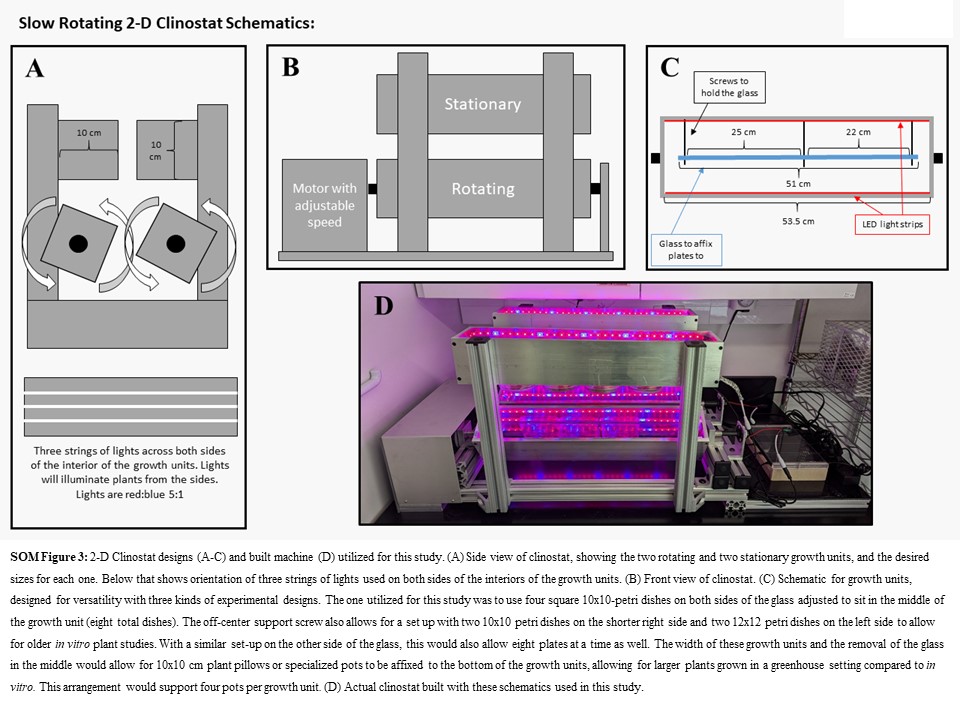
